# Spatial organization of the tumor-immune microenvironment in ER-positive breast cancer: remodeling during treatment and associations with clinical response

**DOI:** 10.1101/2025.10.09.681336

**Authors:** Maria Aanesland Dahle, Jan-Lukas Førde, Eivind Valen Egeland, Allison L. Creason, Cameron Watson, Øystein Garred, Lina Prasmickaite, Gunhild M Mælandsmo, Gordon B. Mills, Olav Engebraaten, Mads Haugland Haugen

## Abstract

**Background:** The tumor microenvironment influences treatment response in ER-positive breast cancer, but what distinguishes responders from non-responders and how it changes during treatment is poorly understood.

**Methods:** ER-positive breast tumors treated with neoadjuvant chemotherapy with or without bevacizumab were profiled with bulk proteomics pre-(n = 95), on-(n = 84) and post-treatment (n = 100). A subset of tumors was profiled with spatial single-cell proteomics pre-(n = 13) and on-treatment (n = 11). Cell phenotypes, spatial location and activation states were determined, and cellular colocalization assessed with spatial metrics. Bulk and spatial features were evaluated against treatment response defined by residual cancer burden.

**Results:** Treatment with bevacizumab amplified chemotherapy effects on proteomic signaling. The immune contexture shifted from suppressive to supportive during treatment through decreased macrophage, regulatory and anergic T-cell density and increased colocalization between epithelial cells and CD8+, CD4+ T-cells and dendritic cells. At baseline, responders had high density of effector memory T-cells, while non-responders had more naïve T-cells. In addition, responders had increased colocalization of epithelial cells with macrophages, and effector memory T-cells with M1-like macrophages compared to non-responders.

**Conclusions:** Spatially distinct tumor-immune microenvironments influence response to neoadjuvant treatment, offering valuable insights for guiding treatment decisions.

## Background

Up to 80% of all breast cancers are estrogen receptor (ER) positive [1], and such patients with locally advanced disease are commonly treated with neoadjuvant chemotherapy (NACT). The NeoAva clinical trial investigated efficacy of NACT combined with anti-angiogenic treatment using bevacizumab (bev+NACT) in HER2-breast cancer patients. The addition of bevacizumab significantly increased pathological complete response (pCR) in ER-positive tumors but not in patients with triple negative breast cancer (TNBC) [2]. Other clinical trials have shown increased pCR rate in HER2-breast cancer with bev+NACT but not increased overall survival [3,4]. A lack of biomarkers able to identify patients likely to benefit contributes to the limited use of bev+NACT in these patients.

Conventional cancer therapies target the tumor cells directly. Intrinsic cancer cell heterogeneity and the tumor microenvironment (TME) – comprising cells with pro- and anti-tumor properties – can influence the efficacy of these classical therapies [5]. In response to this challenge, therapies targeting TME components have been developed. This includes bevacizumab, targeting angiogenesis through inhibition of VEGF signaling [6–8]. In addition to its angiogenic activity, VEGF is believed to facilitate immune evasion and promote a pro-tumor microenvironment through recruitment of T-regulatory cells, enhance the immune suppressive functions of macrophages, and limit T-cell infiltration [9–11]. Vascular normalization induced by anti-angiogenic therapy may convert the immune suppressive environment to an immune supportive [12]. Previous studies of the NeoAva patient cohort have shown that an immune responsive phenotype at baseline is associated with treatment response [13]. A molecular signature, including immune- and TME-related proteins, was demonstrated to predict response to bev+NACT [14]. Despite these findings, the underlying biological mechanisms affecting the TME and improving treatment response remains largely unknown.

Recent advances in multiplex protein imaging technologies have generated vast spatial datasets which allow for detailed single-cell phenotyping and study of tumor-immune cell interactions [15–17]. In breast cancer, tumor infiltrating lymphocytes (TILs) have been shown to provide prognostic and possible predictive information [18], making TNBC patients candidates for immunotherapy [19]. Spatial analysis has defined distinct tumor-immune microenvironments of TNBC, often described as cold or hot, and has been linked to survival [20,21] and response to immunotherapy [22]. Yet, ER-positive breast cancer is traditionally considered a cold subtype, and the established biomarkers mainly concern TNBC or HER2+ breast cancer. Bulk and spatial proteomic profiling of time-series biopsies of ER-positive tumors could provide new insights into the prognostic and predictive role of tumor-intrinsic properties and TME subtypes of these tumors.

In this study, we have investigated spatiotemporal tumor characteristics in ER-positive breast cancers treated with bev+NACT. To characterize changes in tumor-intrinsic properties during therapy and find associations with response, we profiled tumors at three timepoints with bulk proteomics using reverse phase protein array (RPPA). A subset of the tumors was further profiled with cyclic immunofluorescence (cycIF) to investigate how the spatial structure of the tumor-immune microenvironment changes during bev+NACT and influences treatment response.

## Materials and Methods

### Patient cohort

In the neoadjuvant NeoAva (NCT00773695) clinical trial, 132 ER-positive or TNBC patients with large or locally advanced tumors were randomized 1:1 to receive chemotherapy with or without bevacizumab (bev+NACT vs. NACT). All patients received four cycles of FEC100 (5-fluorouracil, epirubicin and cyclophosphamide) every 3 weeks for 12 weeks, followed by docetaxel every 3 weeks or weekly paclitaxel for 12 weeks. Samples included and analyzed in this study were collected from 111 ER-positive patients, in which 57 were included in the NACT arm and 54 in the bev+NACT arm. Breast tumor tissue was collected at baseline, on-treatment (W12) and at time of surgery (W25). The primary endpoint in NeoAva was pathological complete response (pCR) measured at time of surgery. In addition, the response measure residual cancer burden (RCB) was used as response measure, which integrates tumor size, cellularity and nodal burden and found to be an independent prognostic factor in breast cancer [23].

### Bulk proteomics with RPPA

Expression of 210 breast-cancer related proteins was quantified in bulk tumor lysates from all three timepoints using RPPA at the MD Anderson RPPA core facility (Houston, TX, USA). In brief, lysates were serially diluted in 5 two-fold dilutions with lysis buffer and printed on nitrocellulose-coated slides, then probed with 210 validated primary antibodies followed by detection with biotinylated secondary antibodies. The relative protein levels for each sample were determined and normalized for protein loading, then log_2_-transformed and median-centered over proteins. Differential expression of proteins was determined with the R package *limma* with the function *lmFit* and *eBayes*. Pathway activity scores (PAS) of protein signaling pathways calculated for each sample were determined by summarizing protein levels of all proteins positively associated with the pathway and subtracting the protein levels of all proteins negatively associated with the pathway, as previously described [24,25].

### Single-cell spatial proteomics with cycIF

Spatial single-cell protein expression was determined by cycIF imaging in 5 µm tumor tissue FFPE section from a selection of patients in the bev+NACT treatment arm. Tissue samples were sequentially imaged in cycles using four target antibodies and nuclear stain with DAPI in each round (Supplementary Table S1) as previously described [15]. Tissue autofluorescence from each channel was captured in the first cycle.

The images were processed in cancer.usegalaxy.org [26] using the MCMICRO pipeline [27]. From raw czi-images, background and shading correction was performed with the BaSiC tool [28]. The image cycles were registered using the ASHLAR algorithm [29]. Cell segmentation, finding location and boundaries for each cell, was performed with the Mesmer algorithm [30]. The second and last round of DAPI were used as nuclear markers, and CKs, E-Cadherin and CD45 were used as membrane markers. Integrated single-cell expression of each marker was quantified using MCQUANT and background adjusted with signal-to-background ratio of autofluorescence signal. Quality control of each marker was performed by visual inspection of the staining consistency and signal-to-noise on representative images, resulting in exclusion of CD20 and CD11b. To further ensure data quality and minimize edge effects in the spatial analysis, sample outlines and staining defects were manually masked and cells found outside these areas were excluded. For each cell, the distance to nearest edge was recorded for use in analysis sensitive to edge effects. The final dataset included 27-plex protein expression profiles from ∼1 million single cells across baseline (n = 13) and on-treatment (n = 11) samples.

### CycIF cell phenotyping

Cell phenotyping was performed by hierarchical gating of log1p-transformed lineage defining markers. Gate thresholds were manually set for each sample and marker by inspecting marker intensity density plots and visualization of the images overlayed with the cell type calling. Quality control of the cell phenotyping was performed with the Vitessce tool in Galaxy where the cell type calls were overlayed on top of the images [31]. Unassigned cells were labeled as “unknown” and immune cells not matching any subtypes were labeled as “other immune”.

### Spatial metrics

Cellular density was calculated as cell count divided by tissue area. Tissue area was determined by partitioning the tissue space into 250×250 µm tiles and those with less than 25 cells (< 400 cells/mm^2^) were excluded. The average minimum distance (AMD) and the Morisita index were used to quantify the degree of colocalization between two cell types [32]. The AMD from a reference cell type to a target cell type was calculated using the *nn2* function in R package *RANN*. This score determines the proximity of two cell types in the tissue. The Morisita score was calculated by summing the products of the proportions of two cell types across grid squares in tissue space, divided by the summed squared proportions of each cell type. This score reflects how similarly two cell types are distributed within the tissue. Detailed description of the spatial metrics and macrophage characterization is given in Supplementary Methods.

### Statistical analysis

Univariate logistic regression models were fitted using function *glm* with parameter family = “binomial” in R. Pairwise comparisons were performed for relevant groups as indicated using two-sample t-test or Mann-Whitney U test if data did not meet assumption of normality. Comparisons of changes in time-series data were tested with paired t-test. A p-value < 0.05 was used as cutoff for statistical significance and labeled with asterisk (*) in figures. All calculations were performed in R using R-studio with R version 4.3.1 and figures generated using R package *ggplot2*.

## Results

### Bevacizumab amplifies changes in protein expression during NACT in ER-positive breast tumors

The NeoAva clinical trial investigated the efficacy of bev+NACT vs NACT, demonstrating improved pCR rate in ER-positive breast cancer patients [2]. To investigate protein profiles associated with response and treatment-induced changes, we assessed bulk protein expression at baseline (n = 95), on-treatment (n = 84) and at surgery (n = 100) using RPPA and spatially resolved single-cell expression for a subset at baseline (n= 13) and on-treatment (n = 11) using cycIF (Figure 1a). Comparing on-treatment to baseline revealed 140 proteins significantly differentially expressed (FDR < 0.05) in the bev+NACT arm and 75 proteins in the NACT arm (Figure 1b). Protein expressional changes were more pronounced in patients that received bev+NACT, however, 87% of the proteins were similarly up- or down-regulated in both treatment arms (Supplementary table S2). The top upregulated protein on-treatment in the bev+NACT arm was Caveolin-1, and several proteins involved in DNA damage response were significantly downregulated including PARP1, MSH2, MSH6 and 53BP1.

**Fig. 1:**
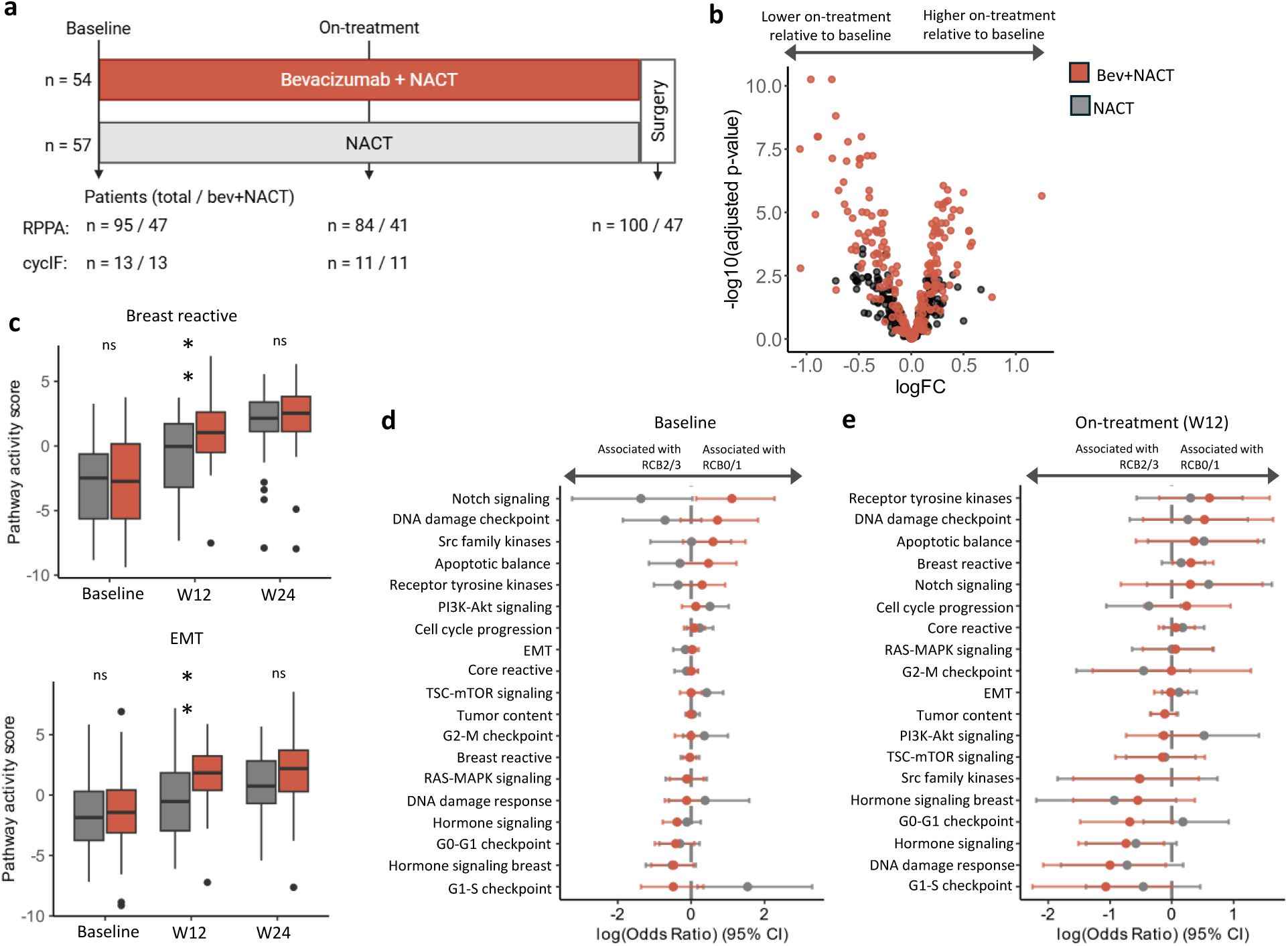
Proteomic response to neoadjuvant chemotherapy ± bevacizumab. **a**, Schematic presentation of the NeoAva clinical trial with longitudinal tumor biopsy sampling and generation of molecular data. **b**, Volcano plot of differentially expressed proteins on-treatment relative to baseline visualized by plotting false discovery rate (FDR) adjusted p-values against the log_2_ fold change in protein expression. **c**, Boxplot of pathway activity scores of the breast reactive pathway and epithelial-mesenchymal transition (EMT) at baseline, on- and post-treatment separated by treatment arm. **d**, Odds ratios for associations between pathway activity scores and RCB 0/1 at baseline, and **e**, on-treatment. Circles represent the point estimate, and whiskers indicate 95% confidence interval. ∗p < 0.05; ∗∗p < 0.01; ns, not significant.

By assessing pathway activity scores of key oncogenic pathways, we found that the tumors changed towards a more reactive breast cancer subtype on-treatment (p < 0.001) (Figure 1c). This subtype is defined by characteristic proteins produced by the microenvironment or cancer-associated fibroblasts such as Caveolin-1, Myosin11, and Rab11 [33]. In line with the protein expressional changes, the reactive breast cancer pathway was significantly amplified in tumors treated with bev+NACT compared to NACT on-treatment (p = 0.006). Similarly, epithelial-mesenchymal transition (EMT) pathway activity, defined by proteins such as Fibronectin and N-Cadherin, was increased on-treatment (p < 0.001) and significantly more active in bev+NACT treated tumors (p = 0.004) (Figure 1c). Protein pathways related to proliferation and cancer progression (G2-M checkpoint, cell cycle progression, tumor content) decreased during treatment and were significantly more suppressed in tumors treated with bev+NACT compared to NACT alone (p < 0.01) (Supplementary Figure S1). In addition, DNA damage response (DDR) was suppressed on-treatment, and significantly more suppressed in tumors treated with bev+NACT (p < 0.01). This pathway is defined by DNA damage repair proteins such as 53BP1, PARP1 and XRCC1, which were also among the top downregulated proteins in the bev+NACT arm. Thus, adding bevacizumab to chemotherapy amplifies changes in protein expression, with stronger suppression of proliferation and DNA repair pathways and increased breast reactive and EMT signaling.

### Tumor-intrinsic protein pathways associated with response to treatment

We investigated whether pathway activity scores at baseline or on-treatment were associated with treatment benefit. In ER-positive breast cancer, achieving RCB 0 or 1 is associated with improved survival and RCB 2 or 3 with worse outcome [34]. Thus, responders were classified as RCB 0/1, and non-responders as RCB 2/3. Univariate logistic regression models were fitted to determine pathway activity scores and associations with RCB 0/1 at baseline (Figure 1d) and on-treatment (Figure 1e). Notably, Notch signaling was the only pathway significantly associated with response to bev+NACT at baseline (p = 0.019) and was negatively associated with response to NACT alone. Low DDR and several pathways linked to cancer progression were negatively associated with response to bev+NACT on-treatment: hormone signaling, G0-G1 checkpoint, hormone receptor and G1-S checkpoint.

### The single-cell spatial proteomic landscape in ER-positive tumors

To further elucidate protein compositional changes during treatment with bev+NACT, a subset of tumors was profiled by single-cell spatial proteomics. Comprehensive TME profiles were captured with a 27-plex antibody panel, particularly of the immune compartment (Figure 2a). A total of 24 multiplex images representing 13 patients at baseline paired with 11 patients on-treatment were acquired. Following image acquisition and single-cell segmentation, cells were phenotyped using a hierarchical gating strategy (Figure 2b). This identified major lineages: epithelial (Cytokeratin (CK+)), stromal (αSMA+), immune (CD45+) and endothelial cells (CD31+, CK-, CD45-). CD45+ immune cells were further sub-classified into CD8+ T cells, CD4+ T cells, regulatory T cells (Tregs), macrophages, and dendritic cells (DCs). This resulted in eight primary cell type classifications, along with ‘Other Immune’ and ‘Unknown’ categories, for detailed spatial analysis (Figure 2c).

**Fig. 2:**
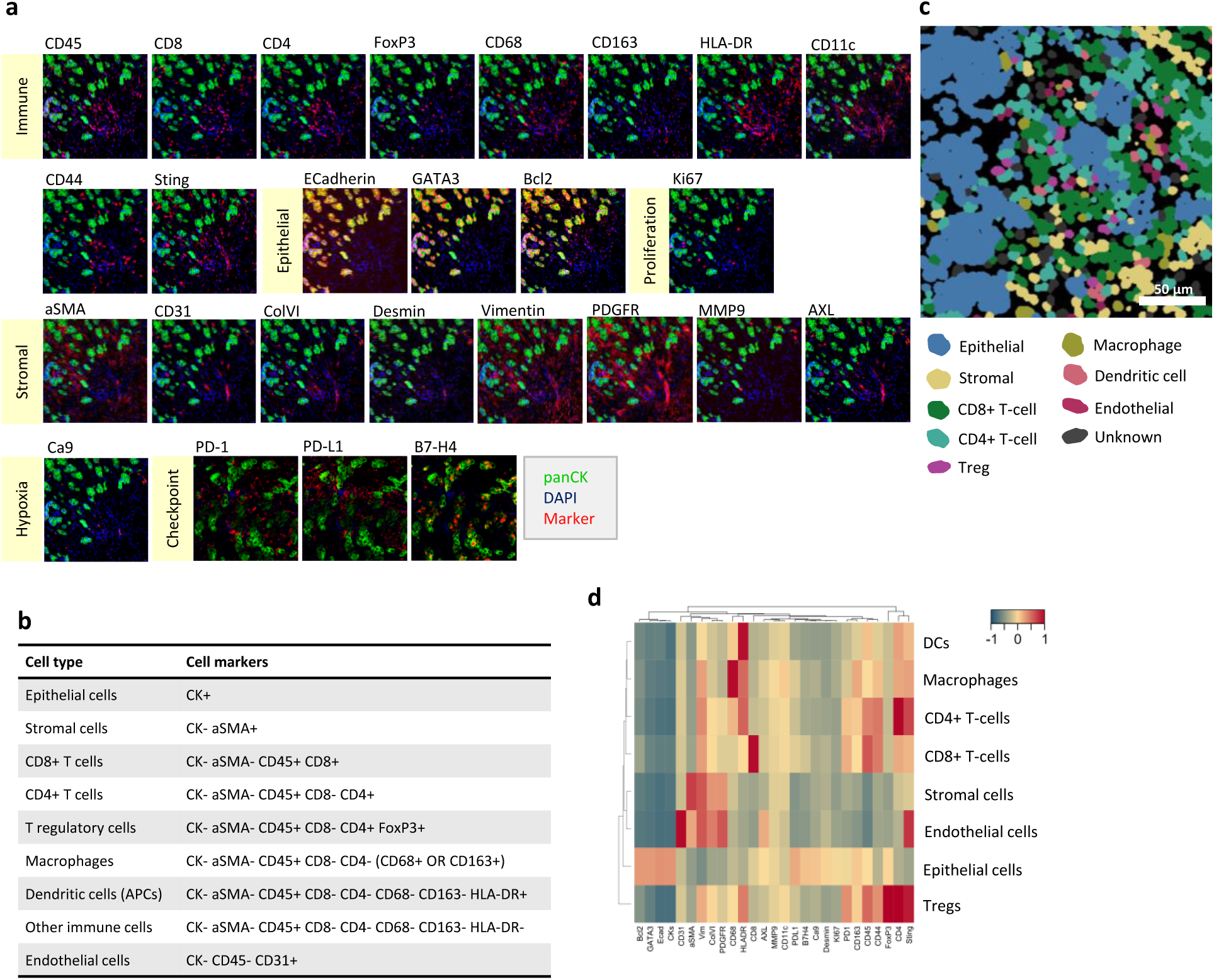
Longitudinal multiplexed imaging with cycIF. **a,** Representative images of protein expression from 27-plex antibody panel. **b**, Hierarchical gating strategy for cell phenotyping. **c**, Representative image illustrating cell segmentation and cell phenotypes. **d**, Heatmap of mean protein expression in eight cell phenotypes grouped with hierarchical clustering.

Epithelial cells were distinguished by high expression of CK, E-cadherin, GATA3 and Bcl2 (Figure 2d). Stromal and endothelial cells were characterized by high expression of aSMA, Vimentin, CollagenVI and PDGFR, while the endothelial cells also expressed high amounts of CD31, Sting and AXL. All immune cells except Tregs were grouped in hierarchical clustering, and were characterized by high expression of CD45, CD44 and Sting in addition to their respective lineage identification markers. Tregs had the highest expression of PD-1 compared to the other cell types, but expression of this marker was also found on CD8+ T-cells and CD4+ T-cells. PD-L1 was expressed at high levels by epithelial cells. The cell types classified as “unknown” and “other immune” were not distinguished by any specific marker and therefore not used in the subsequent analysis (Supplementary Figure S2).

### Cell type composition dynamics during treatment

To assess how the TME changes during therapy, cell type composition and density at baseline and on-treatment was analyzed. The dominating cell types at baseline were epithelial and stromal cells, while endothelial and immune cells were present in higher fractions on-treatment (Figure 3a). Sample S1 was a distinct outlier at baseline, characterized by a remarkably dense immune infiltrate rich in CD8+ T cells. Despite a general decrease in epithelial fraction on-treatment, high heterogeneity was observed in epithelial and immune fractions at both timepoints. Illustrating this, sample S3 presented with high immune cell fractions at both timepoints, contrasting with persistently low immune fractions in S10 (Figure 3b). The overall cell density decreased from 3739 cells/mm^2^ in baseline samples to 1821 cells/mm^2^ on-treatment, driven by a significant decrease in epithelial and stromal cell density (p < 0.001 and p = 0.023, respectively) (Figure 3c). Cell density of CD8+, CD4+ T-cells and DCs remained constant, suggesting that the tumors did not experience an immune cell infiltration in response to treatment. Notably, density of macrophages and Tregs decreased significantly (p = 0.024 and p = 0.023, respectively), demonstrating a reduction in immune suppressive populations on-treatment.

**Fig. 3:**
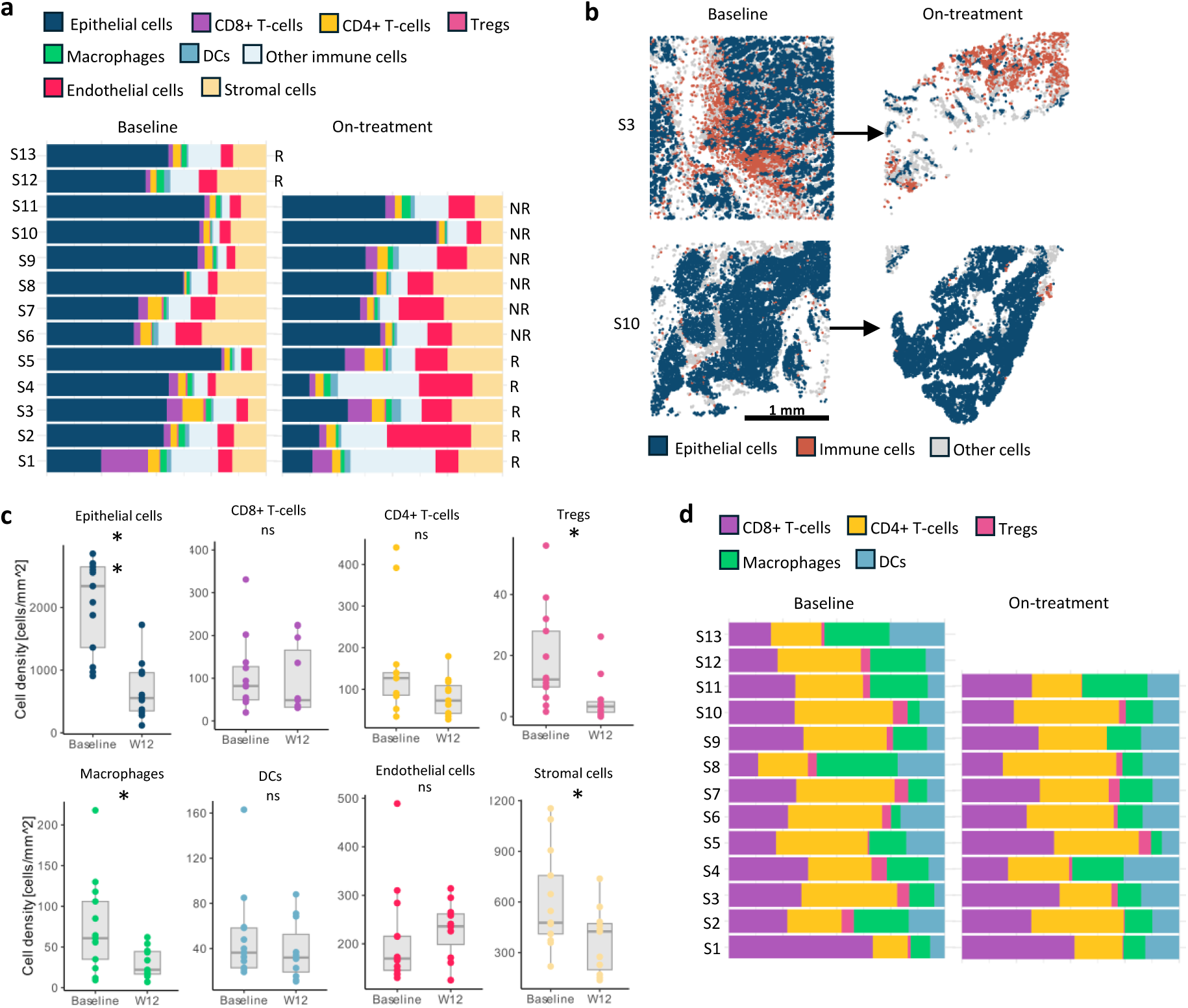
Longitudinal TME cell type composition. **a**, Stacked bar plot of cell type fractions in samples S1-S13 at baseline and on-treatment (R: responder, NR: non-responder). **B**, Representative sections of epithelial (blue) and immune cell (red) spatial location of sample S3 and S10 at baseline and on-treatment. **c**, Boxplot of epithelial, CD8+ T-cell, CD4+ T-cell, Treg, DC, endothelial and stromal cell density at baseline and on-treatment. **d,** Stacked bar plot of immune cell type fractions in samples S1-S13 at baseline and on-treatment. ∗p < 0.05; ∗∗p < 0.0; ns, not significant.

To understand the relative balance of immune populations, the composition within the annotated immune cell compartment was analyzed separately (Figure 3d). The immune compartment was not particularly dominated by either myeloid or T-cell types. At baseline, CD8+ T-cells comprised an average of 30±12 % (mean ± SD) of all annotated immune cells and 33±10 % on-treatment. CD4+ T-cells consisted of 34±10 % at baseline, and 35±10 % on-treatment. Thus, the composition of the major immune cell types remained stable and did not shift from myeloid to lymphoid dominated or vice versa following treatment.

### Spatial remodeling of the TME during treatment

Changes in the tissue architecture of tumors during treatment were explored using two spatial metrics: average minimum distance (AMD), and the Morisita index (Figure 4a). AMD was used to determine cell to cell proximity and is sensitive to variations in cellular density. The Morisita index, further denoted “colocalization score”, was used to quantify how similarly two cell types were distributed in the tissue.

**Fig. 4:**
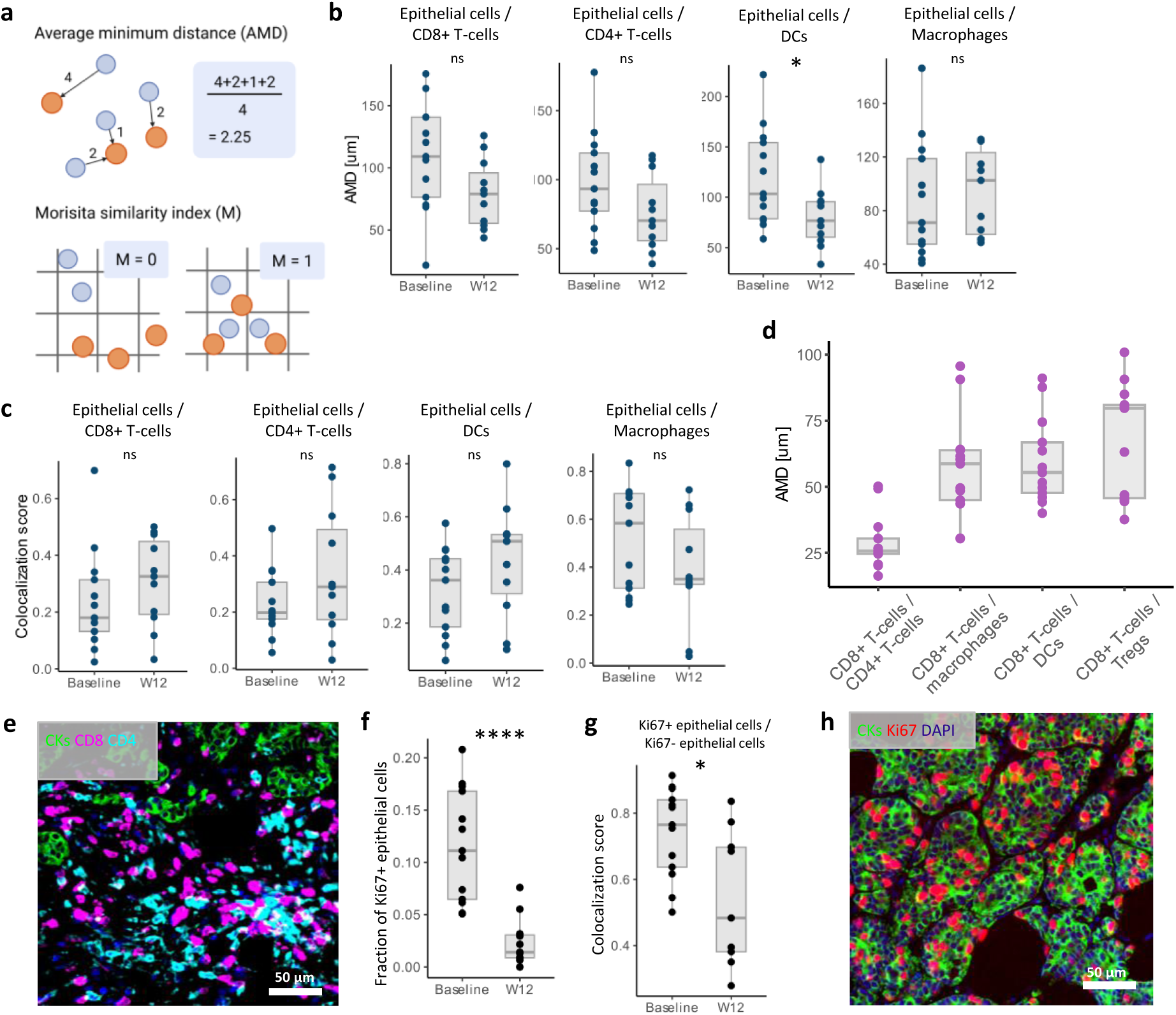
Spatial remodeling of the TME. **a,** Schematic figure illustrating the calculation of AMD and colocalization score (detailed description is given in Supplementary Methods). **b**, Boxplot of AMD between epithelial cells and CD8+, CD4+ T-cells, DCs and macrophages at baseline and on-treatment. **c**, Boxplot of colocalization score between epithelial cells and CD8+, CD4+ T-cells, DCs and macrophages at baseline and on-treatment. **d**, Boxplot of AMD between CD8+ T-cells and CD4+ T-cell, Tregs, DCs and macrophages at baseline. **e**, Representative image of colocalization between CD8+ and CD4+ T-cells. **f**, Boxplot of Ki67+ fraction of epithelial cells at baseline and on-treatment. **g**, Boxplot of colocalization score between Ki67+ and Ki67-epithelial cells. **h**, Representative image of Ki67 expression in epithelial cells. ∗p < 0.05; ∗∗p < 0.01; ****p<0.0001; ns, not significant. AMD: average minimum distance.

The colocalization score is less dependent on cellular density than AMD. We first investigated spatial interactions of epithelial cells and other cell types. At baseline, an epithelial cell was, on average, around 100 µm away from the closest immune cell. Cell to cell proximity revealed that epithelial cells were closer to CD8+ T-cells, CD4+ T-cells and DCs on-treatment compared to baseline (p = 0.05, 0.08, 0.03, respectively), while distance to macrophages and Tregs were stable (Figure 4b, Supplementary Figure S3). The colocalization score revealed that epithelial and immune cells were independently or separately distributed across the tissue at baseline (Figure 4c). In line with the proximity measures, colocalization between epithelial cells and CD8+ T-cells, CD4+ T-cells and DCs increased during treatment. Taken together, these results could suggest more spatial interactions between epithelial cells and anti-tumor immune cells on-treatment, and remodeling from an immune suppressive towards an immune supportive TME during treatment.

Given the importance of CD8+ T-cells in anti-tumor immunity, spatial interactions with other immune cells were explored. At baseline, CD8+ T-cells were found near CD4+ T-cells, while distance to Tregs, DCs and macrophages was generally greater (Figure 4d, e). Consistently high colocalization score and low AMD between CD8+ and CD4+ T-cells suggest that these cells are commonly found in close proximity within the TME, while CD8+ T-cells and Tregs, DCs and macrophages were spatially independent of one another (Supplementary Figure S4). The spatial immune organization was persistent on-treatment, and no significant changes in colocalization pattern were observed.

Since chemotherapy targets proliferating cancer cells, changes in proliferation state of epithelial cells during therapy were investigated. As expected, the fraction of Ki67+ epithelial cells significantly decreased from baseline to on-treatment (p < 0.001) (Figure 4f). At baseline, Ki67+ and Ki67-epithelial cells were highly colocalized, indicating that proliferating and non-proliferating epithelial cells were similarly distributed across the samples (Figure 4g, h). Colocalization decreased during treatment, which could indicate, together with a low Ki67+ fraction, a shift toward predominantly non-proliferative epithelial regions on-treatment.

### Macrophage density and colocalization with epithelial cells are associated with clinical response

We further investigated whether the tissue architecture of baseline and on-treatment tumors was linked to treatment response. At baseline, there was no difference in epithelial cell density in responders and non-responders. As the density decreased on-treatment, the epithelial cell density was significantly lower in responders compared to non-responders (p = 0.045), showing that treatment effects can be observed at the cellular level 12 weeks after start of treatment (Figure 5a). Notably, high density of macrophages at baseline was significantly associated with response (p = 0.022) (Figure 5a). No other significant differences in cellular density between responders and non-responders were observed at either timepoints (Supplementary Figure S5).

**Fig. 5:**
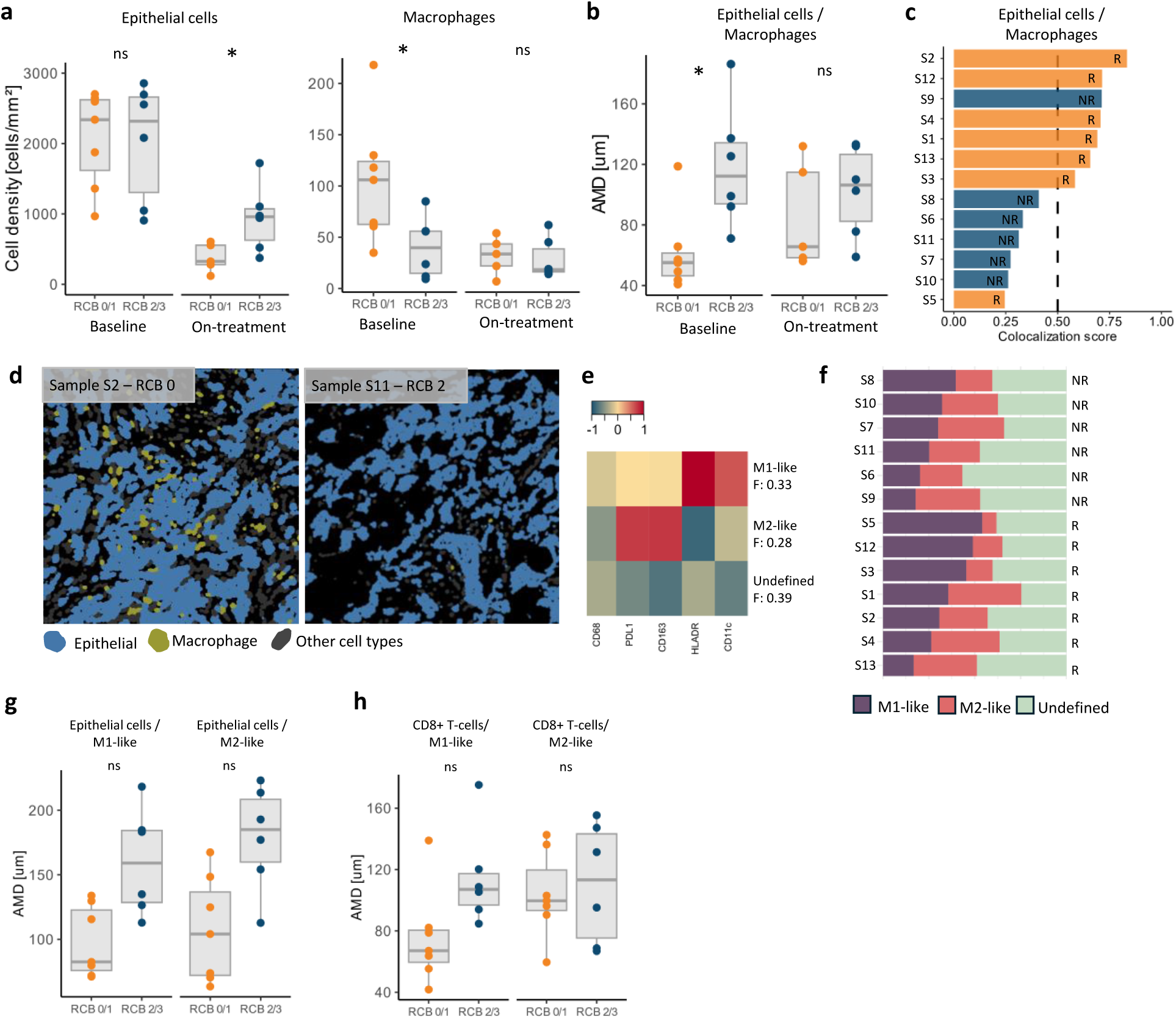
Spatial features associated with response to treatment. **a**, Boxplot of epithelial cell and macrophage density in responders (n = 7 baseline, n = 5 on-treatment) and non-responders (n = 6 baseline, n = 6 on-treatment) at baseline and on-treatment. **b**, Boxplot of AMD between epithelial cells and macrophages separated by responders and non-responders at baseline and on-treatment. **c**, Colocalization score between epithelial cells and macrophages at baseline sorted from highest to lowest score (R: responders; NR: non-responders). **d**, Representative section of epithelial cell and macrophage spatial location in sample S2 and S11. **e**, Heatmap of mean protein expression in three macrophage phenotypes (f: fraction). **f**, Stacked bar plot of macrophage subtype fraction at baseline in samples S1-S13 separated by responders (R) and non-responders (NR) and sorted from lowest to highest M1-like fraction. **g**, Boxplot of AMD between epithelial cells and M1-like and M2-like macrophages. **h**, Boxplot of AMD between CD8+ T-cells and M1-like and M2-like macrophages. ∗p < 0.05; ∗∗p < 0.01; ns, not significant. AMD: Average minimum distance.

Next, the association between cellular colocalization and treatment response was examined. Notably, epithelial cells and macrophages were highly colocalized in responders at baseline. In responders, epithelial cells were located ∼60 µm from the closest macrophage on average (Figure 5b). Non-responders had an average colocalization score of 0.38 compared to 0.63 for responders at baseline, meaning that epithelial cells and macrophages were colocalized in responders and spatially separated in non-responders (Figure 5c). There was no significant difference in colocalization of epithelial cells and other evaluated cell types in responders and non-responders at either timepoint (Supplementary Figure S6). Altogether, this indicated that increased abundance of macrophages and their proximity to tumor cells at baseline correlated with better response to treatment (Figure 5d).

### Spatial organization of macrophage subtypes in the TME

Given the dual role of macrophages in the TME, classically described as M1-like (anti-tumor) and M2-like (pro-tumor), the macrophage population at baseline was further characterized by hierarchical clustering using relevant markers: CD163, PD-L1, HLA-DR and CD11c. This resulted in three macrophage clusters: one pro-inflammatory (M1-like) expressing high levels of HLA-DR and CD11c, one immune suppressive (M2-like) expressing high levels of CD163 and PD-L1, and one undefined with low expression of subtype markers (Figure 5e). Although the macrophage composition varied between the samples, there was no significant difference between responders and non-responders (Figure 5f).

We next examined colocalization of epithelial cells and CD8+ T-cells with the macrophage subtypes in responders and non-responders. Our findings showed that in responders, epithelial cells were in closer proximity to both M1- and M2-like macrophages compared to non-responders (Figure 5g). Furthermore, CD8+ T-cells were found to be closer to M1-like macrophages in responders, suggesting potential interactions between these cell types (Figure 5h).

### Distinct CD8+ T-cell subtypes are associated with clinical response

Since CD8+ T-cells and M1-like macrophages were closely interacting in responding patients, the CD8+ T-cell population was further characterized. Using CD44 to identify memory T-cells and PD-1 to depict sustained antigen exposure, four T-cell populations were defined: naïve T-cells (T_naïve_, CD44-/PD-1-), anergic naïve T-cells (T_AN_, CD44-/PD-1+), central memory T-cells (T_CM_, CD44+/PD-1-), and effector memory T-cells (T_EM_, CD44+/PD-1+) (Figure 6a). Association with clinical response was explored in these subpopulations. Notably, responders had significantly higher fraction and density of T_EM_-cells (p = 0.002 and p = 0.004, respectively) as well as T_CM_-cells and T_AN_-cells at baseline (Figure 6b, Supplementary Figure S7). Non-responders had significantly higher fraction, but not density, of T_naïve_ -cells (p = 0.008). This indicates that responders had high diversity of T-cell populations enriched for potential tumor reactive subtypes while non-responders mainly had high levels of T_naïve_-cells. In addition, proliferating T_EM_-cells were identified using Ki67, demonstrating significantly higher fraction and density of Ki67+ T_EM_-cells in responders (p = 0.004 and p = 0.003, respectively) (Figure 6c, Supplementary Figure S7).

**Fig. 6:**
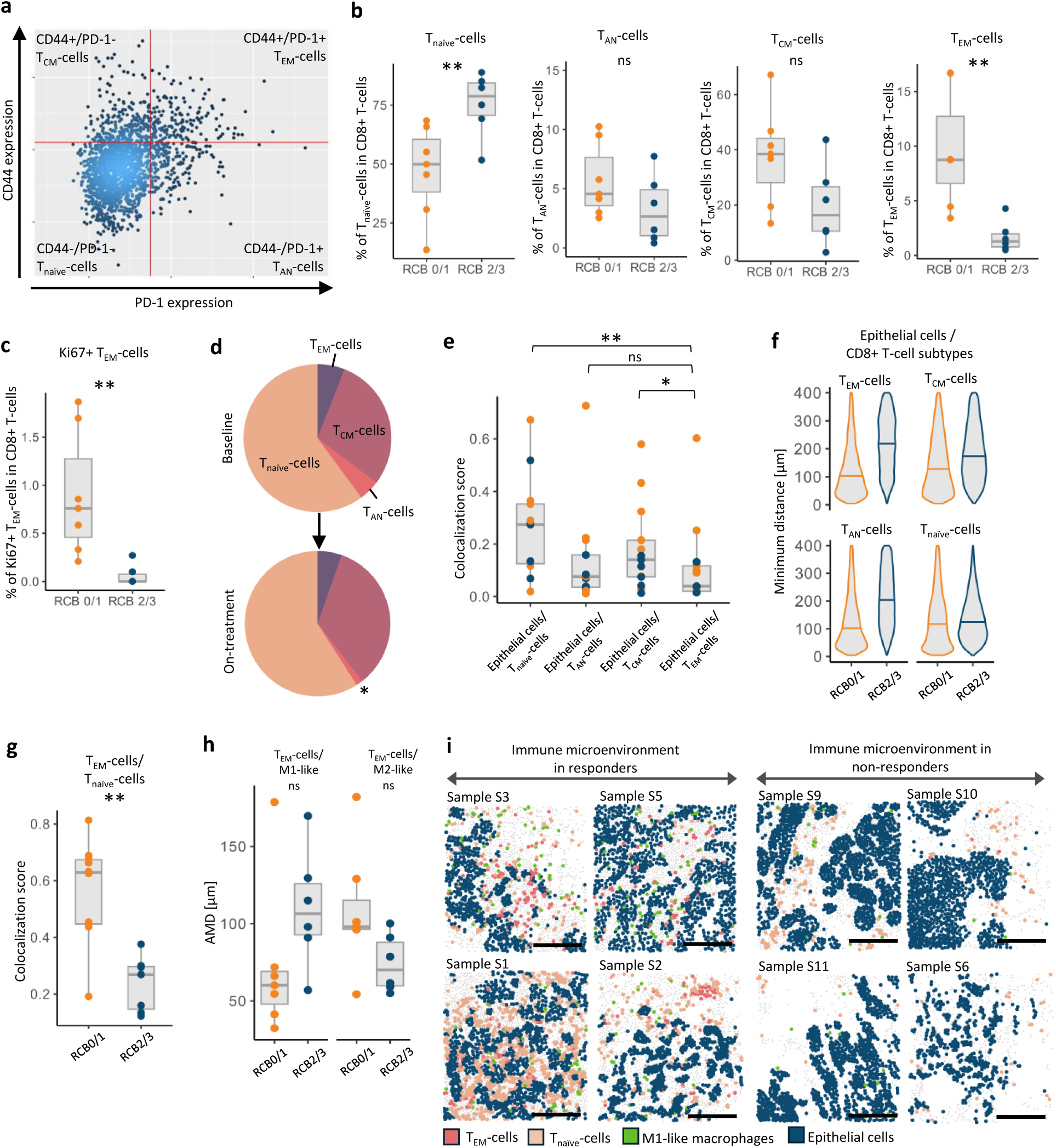
Distinct CD8+ T-cell populations and colocalization is linked to response. **a,** Representative CD44 and PD-1 expression in CD8+ T-cells at baseline. **b**, Percentage of CD8+ T-cell subtypes at baseline separated by responders and non-responders. **c**, Percentage of Ki67+ T_EM_-cells in CD8+ T-cells separated by responders and non-responder. **d**, Pie chart of average CD8+ T-cell subtype fraction at baseline and on-treatment. **e**, Colocalization score of epithelial cells and CD8+ T-cell subtypes. **f**, Violin plot of single-cell minimum distance between epithelial cells and CD8+ T-cell subtypes separated by responders and non-responders. **g**, Colocalization score between T_EM_-cells and T_naïve_-cells separated by responders and non-responders. **h**, AMD between T_EM_-cells and M1-like and M2-like macrophages separated by responders and non-responders. **i**, Sections of T_EM_-cell, T_naïve_-cell, M1-like macrophage, and epithelial cell spatial location in representative samples. Scalebar: 250 µm. ∗p < 0.05; ∗∗p < 0.01; ns, not significant. AMD: average minimum distance.

Overall, the fraction of T_EM_-cells and T_naïve_-cells was stable from baseline to on-treatment, while the fraction of T_AN_-cells decreased significantly (p = 0.038) (Figure 6d, Supplementary Figure S8). These results suggest that the level of immune suppressive T-cells decreased and are in line with our previous findings on cellular composition and colocalization during treatment showing that the tumors were changing towards a more immune supportive TME following treatment.

### Effector memory T-cells are highly colocalized with naïve T-cells and M1-like macrophages in responders

The spatial architecture of CD8+ T-cell subtypes was investigated by colocalization and cell to cell proximity measures. In general, epithelial cells were significantly more colocalized with T_CM_-cells and T_naïve_-cells compared to T_EM_-cells, revealing that epithelial cells were closer to PD-1-negative T-cells than PD-1-positive T-cells (Figure 6e). Comparing responders to non-responders revealed that epithelial cells were closer to T_EM_-cells, T_CM_-cells and T_AN_-cells in responders, suggesting closer interactions between these cell types (Figure 6f). In addition, T_EM_-cells were highly colocalized with T_naïve_-cells (p < 0.01) and M1-like macrophages (p = 0.10), while spatially separated to M2-like macrophages (p = 0.10) in responders compared to non-responders (Figure 6g, h). These results show that the immune microenvironment in responders is spatially distinct from non-responders and the immune contexture at baseline influences treatment response (Figure 6i).

## Discussion

In this study, we characterized the effects of adding bevacizumab to NACT on protein expression as well as TME composition and architecture using paired time-series breast cancer biopsies. By profiling ER-positive breast tumors with bulk and single-cell spatial proteomics we found that adding bevacizumab to NACT amplifies tumor-intrinsic proteomic responses and reshapes the immune contexture. To our knowledge, this is the first time ER-positive breast tumors treated with bev+NACT have been profiled with single-cell spatial proteomics.

Protein expressional changes revealed that breast reactive signaling increased on-treatment and was significantly more pronounced in tumors treated with bev+NACT. Previous proteomic studies have identified reactive breast cancer as an independent subtype, consisting mainly of ER-positive tumors, only detected at the protein level [33,35]. This subtype is characterized by high expression of stromal proteins such as Caveollin-1 and Collagen-VI and shown to have a high fibroblast content reflected by high EMT signaling [36]. With spatial analysis, we reported a decrease in stromal cells and an increase in endothelial cells on-treatment. This discrepancy could suggest that it is not the identified stromal cells, but rather the endothelial cells that represent the increased reactive signaling observed with RPPA. Increased stromal signaling together with the reported increase in endothelial cells might indicate vascular normalization on-treatment. This has previously been observed after treatment with bevacizumab and linked to improved clinical response [37,38]. Consistent with this, signaling pathways related to proliferation decreased significantly more in tumors treated with bev+NACT compared to NACT alone, supporting the hypothesis that bevacizumab enhances chemotherapy delivery through vascular normalization. We also observed that tumor-intrinsic signaling pathways related to cancer progression were negatively associated with response on-treatment, and significantly lower density of epithelial cells was observed in responders. These findings are in line with previous studies on the same patient cohort showing that RNA disruption, associated with tumor cell death, mid treatment predicted treatment response [39].

In addition to stromal and tumor-intrinsic changes during treatment, we reported remodeling of the immune microenvironment. Density of Tregs, macrophages and T_AN_-cells decreased significantly, while colocalization of epithelial cells with CD8+ T-cells, CD4+ T-cells and DCs increased simultaneously. This demonstrates that the immune suppressive TME shifted towards an immune supportive state during treatment. These findings align with similar observations that bevacizumab normalizes the tumor vasculature and at the same time reprograms the tumor immune microenvironment [40,41]. Given that such modulation is critical for response to immune checkpoint inhibitors, our results support the use of bevacizumab in combination with immunotherapy and NACT in ER-positive breast cancer. This has previously demonstrated promising clinical benefit in TNBC [42].

Unexpectedly, we found that macrophage density, rather than CD8+ and CD4+ T-cell density, was positively associated with treatment response. This finding challenges the assumption that tumor infiltrating lymphocytes (TILs) as part of the TME are the major contributor to treatment response and good prognosis, which has been shown in TNBC and HER2+ breast cancer [43–47]. The role of TILs in ER-positive breast cancer remains unclear, and our findings illustrate the importance of qualitative and quantitative differences of T-cell subsets. High fraction of T_EM_-cells and low fraction of T_naïve_-cells were associated with response to treatment. Although high levels of PD-1-positive T-cells are linked to T-cell exhaustion and an immune suppressive environment, studies in ER-positive breast cancer have also reported that high levels at baseline were associated with good prognosis [48,49]. Some studies show that PD-1-positive T-cells may resemble tissue resident memory T-cells rather than exhausted T-cells and could provide local immune protection against tumor rechallenge [50,51]. Further, the presence of PD-1-positive T-cells may represent response to neoantigens on the tumor cells. Most likely, the identified T_EM_-cell population is heterogeneous and may also include exhausted T-cells. However, enrichment of proliferating T_EM_-cells in responders indicates that many of these cells are actively cycling and responding to antigen stimulation. High colocalization of T_EM_-cells and T_naïve_-cells in responders was observed, while these T-cell subtypes were spatially separated in non-responders. Spatial proximity could, in addition to displaying cellular interactions, also reflect phenotypic plasticity. These results could therefore indicate that the CD8+ T-cells in responders are, after encountering tumor antigens, in transition from naïve to effector memory to exhausted. This transition, and its association with tumor reactivity, has previously been observed in melanoma [52].

We showed that high density of macrophages in close proximity to epithelial cells was associated with response to treatment. In line with our findings, a prior study showed that high levels of tumor associated macrophages (TAMs) and their proximity to tumor cells were associated with good clinical outcome in gastric cancer [53]. The canonical marker for macrophages, CD68, has previously been shown to have antigen presenting functions [54]. Therefore, macrophages near epithelial cells could reflect an enhanced antigen presentation in these tumors.

Macrophages have dual roles in the TME and can have both M1- and M2-features, acting pro- and anti-tumorigenic depending on where they are on the spectrum [55]. The macrophage subtype composition was not indicative of treatment response in this study, but spatial organization revealed important differences between responders and non-responders. Responders displayed spatial proximity of T_EM_-cells and M1-like macrophages, and non-responders displayed spatial proximity of T_EM_-cells and M2-like macrophages further away from epithelial cells. These results show that distinct immune microenvironments could influence treatment response by enabling effective antigen presentation or production of proinflammatory cytokines by M1-like macrophages to CD8+ T-cells, and promote their transition from T_naïve_-cells to T_EM_-cells. In line with this, high levels of proinflammatory macrophages have previously been observed in TNBC subtypes with high CD8+ T-cell infiltration [21]. In addition, a previous study showed that TAMs, rather than tumor cells, are the primary source of antigen exposure governing transition of progenitor exhausted T-cells to terminally exhausted in glioblastoma [56]. Our results shed light on TAMs complex role in the TME and the importance of investigating cells in the immune contexture beyond TILs.

There are several strengths and limitations to our study. It is difficult to capture the full complexity of the breast cancer disease, known for its heterogeneity, with small tissue specimens. Samples with small fields of view (∼ 1 mm^2^) or TMA cores have been shown to be too small to accurately determine simple features such as cell composition because the sample is too small relative to its feature size. In this study we have used whole-slide images, capturing an average of ∼10^5^ cells at baseline, which have been found to largely overcome this problem [57]. The on-treatment samples were substantially smaller, typically ∼10^4^ cells, and caution is therefore required when drawing conclusions. Another limitation is that sensitivity and specificity of antibodies could influence cell phenotyping. We did conservative manual phenotyping of key lineages to identify cell types, compared to unsupervised clustering. This method gives reliable identification of biologically established phenotypes, with the drawback that a group of cells had to be classified as “unknowns”. Secondly, our marker panel precluded inclusion of some key immune cells, such as B-cells and NK cells. Thus, a complete overview of the immune composition in the tumors could not be given. Overall, the number of samples analyzed makes our findings exploratory and need to be confirmed in larger cohorts.

In conclusion, we combined bulk and single-cell spatial proteomics to map TME dynamics of ER-positive breast cancer treated with bev+NACT. Addition of bevacizumab amplified chemotherapy-induced protein signaling and importantly remodeled the TME from immune-suppressive towards a more immune-supportive state. Treatment response was associated with a pro-inflammatory TME at baseline, where macrophage-tumor proximity and enrichment of T_EM_-cells outperformed TILs as predictors. Spatially distinct immune architectures in responders and non-responders may advance current stratification of ER-positive breast cancer patients and guide treatment decisions in the future.

## Additional Information

## Acknowledgements

The Functional Proteomics Reverse Phase Protein Array Core was supported in part by The University of Texas MD Anderson Cancer Center, P30CA016672 and R50CA221675.

The authors would like to thank the Oregon Health & Science University (OHSU) Knight Cancer Institute Precision Oncology Program, in particular Hongli Ma, David Kilburn and Dong Zhang, for their support and contributions to the cycIF analysis conducted in this study. CycIF imaging at OHSU was supported in part by Advanced light microscopy core in P30CA069533, the Breast Cancer Research Foundation, and a kind gift from the Dr. Miriam and Sheldon Adelson Medical Research Foundation and NCI U01 CA281902 to GBM.

The authors would like to thank Jeremy Goecks for generous access to computational resources in cancer.usegalaxy.org.

Preliminary results of this manuscript were presented at the European Association for Cancer Research (EACR) congress of 2025 [58].

## Authors’ contributions

MAD, JLF and MHH drafted the manuscript. MAD and MHH performed bulk proteomic RPPA analysis. MAD, GBM and MHH coordinated cycIF imaging. MAD, EVE, AC, CW and MHH processed cycIF images. MAD performed cell phenotyping. MAD and JLF performed spatial statistical analysis. GMM, GBM, OE and MHH conceived and secured funding for the study. MHH supervised the work. All authors contributed to data interpretation, manuscript revision and approved the final manuscript.

## Ethics approval and consent to participate

The study was performed in accordance with the Declaration of Helsinki. The NeoAva clinical trial (NCT00773695) was approved by the Regional Ethics Committee for Medical and Health Research Ethics South-East Norway (REK, #2010/335). Informed consent was obtained from all participants involved in the study.

## Data availability

Normalized RPPA data from three timepoints are available as Supplementary Table S3. Raw cycIF images are available upon reasonable request. Spatial single-cell data including identified major cell lineages are available on Zenodo data repository (https://doi.org/10.5281/zenodo.17279174).

## Competing interests

MHH, OE, GBM and GMM have a patent application (PCT/EP2021/052016) related to the bulk proteomic RPPA data. OE receives institutional support from AstraZeneca. GBM is a SAB member or consultant for Amphista, Astex, AstraZeneca, BlueDot, Ellipses Pharmaceuticals, ImmunoMET, Leapfrog Bio, Bruker/Nanostring, Neophore, Nerviano, Nuvectis, Pangea, PDX Pharmaceuticals, Qureator, Rybodyne, Signalchem Lifesciences, Turbine, Zentalis Pharmaceuticals. GBM has stock/options/financial relationships with Bluedot, Catena Pharmaceuticals, ImmunoMet, Nuvectis, RyboDyne, SignalChem Lifesciences, Turbine. GBM has licensed technology: HRD assay to Myriad Genetics, DSP patents with Nanostring. GBM has sponsored research from AstraZeneca, Zentalis, Nanostring and Nerviano. The authors declare that the competing interests have not influenced the content of this manuscript. The authors declare no other conflicts of interest.

## Funding information

MAD, JLF, GMM and MHH were funded by the South-East Norway Regional Health Authorities, Norway (projects #2020072 and #2024099 to MHH, #2021049 to GMM). GMM and OE received funding from the Norwegian Cancer Society (#190257 to GMM and #198091 to OE). RESCUER project has received funding from the European Union’s Horizon 2020 Research and Innovation Program under Grant agreement # 847912. CW and AC had support through the Knight Cancer Institute SMMART Clinical Trials Program and National Institutes of Health/National Cancer Institute (NIH/NCI; U24CA284167).

## Notes

### Competing Interest Statement

The authors declare the following competing interests: Mads Haugland Haugen, Olav Engebraaten, Gunhild M Maelandsmo and Gordon B. Mills have an international patent application (PCT/EP2021/052016) related to the bulk proteomic data used in the study. Olav Engebraaten receives institutional support from AstraZeneca. Gordon B. Mills has stock/options/financial relationships with Bluedot, Catena Pharmaceuticals, ImmunoMet, Nuvectis, RyboDyne, SignalChem Lifesciences, Turbine. Gordon B. Mills has licensed technology: HRD assay to Myriad Genetics, DSP patents with Nanostring. Gordon B. Mills has sponsored research from AstraZeneca, Zentalis, Nanostring and Nerviano. The authors declare that competing interests have not influenced the content of this manuscript and declare no other conflicts of interest.

### Summary of Updates

PDF size changed to A4 format for the figures. Competing interest updated.

