## Supplementary Figures for "Spatial organization of the tumor-immune microenvironment in ER-positive breast cancer: remodeling during treatment and associations with clinical response"

**Supplementary Fig. S1: Longitudinal pathway activity scores**

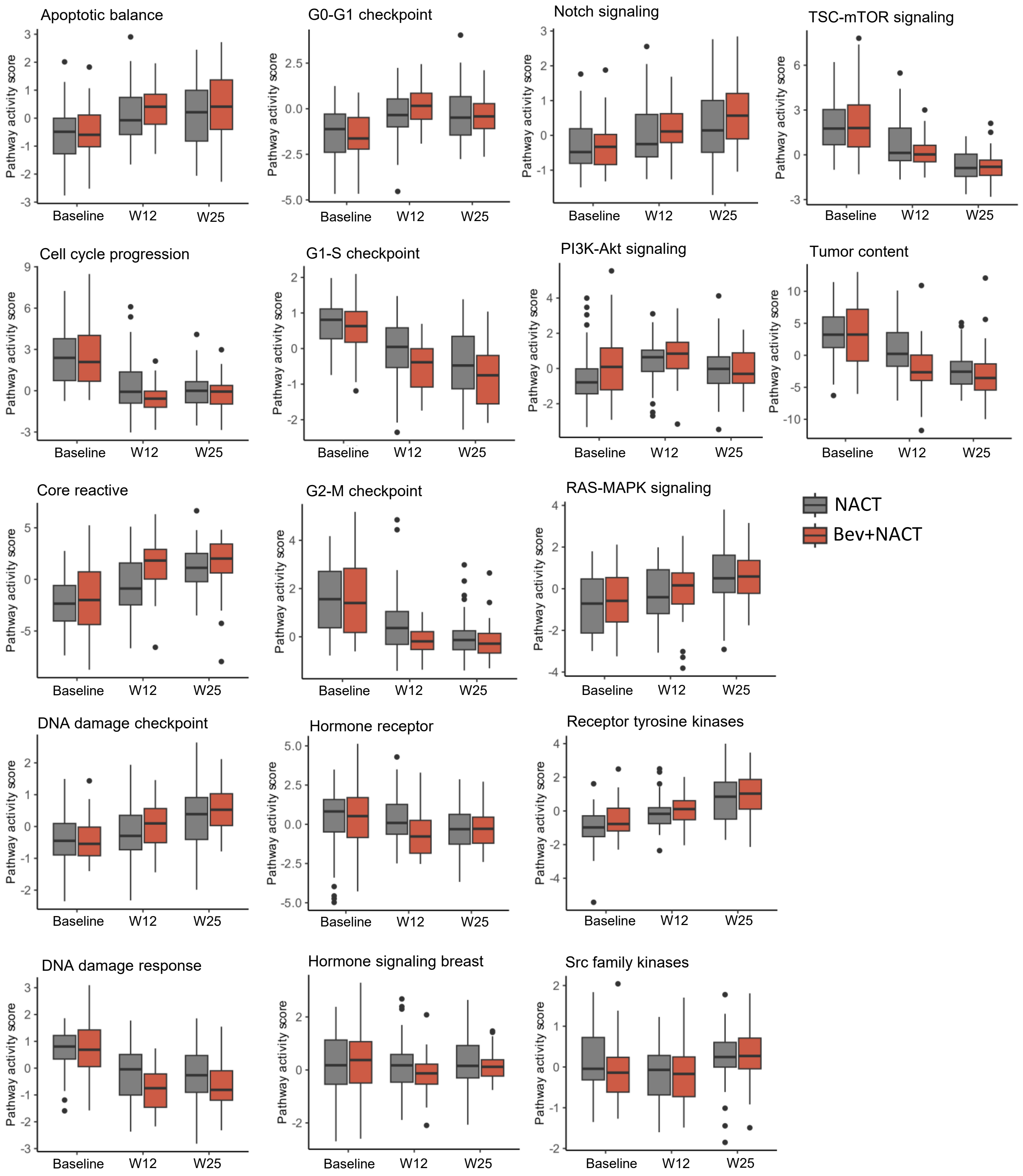


**Supplementary Figure S1. Longitudinal pathway activity scores**. Boxplot of pathway activity scores at baseline, on-treatment (W12) and post-treatment (W24) separated by treatment arm (bev+NACT vs. NACT).

**Supplementary Fig. S2: Cell type protein expression**

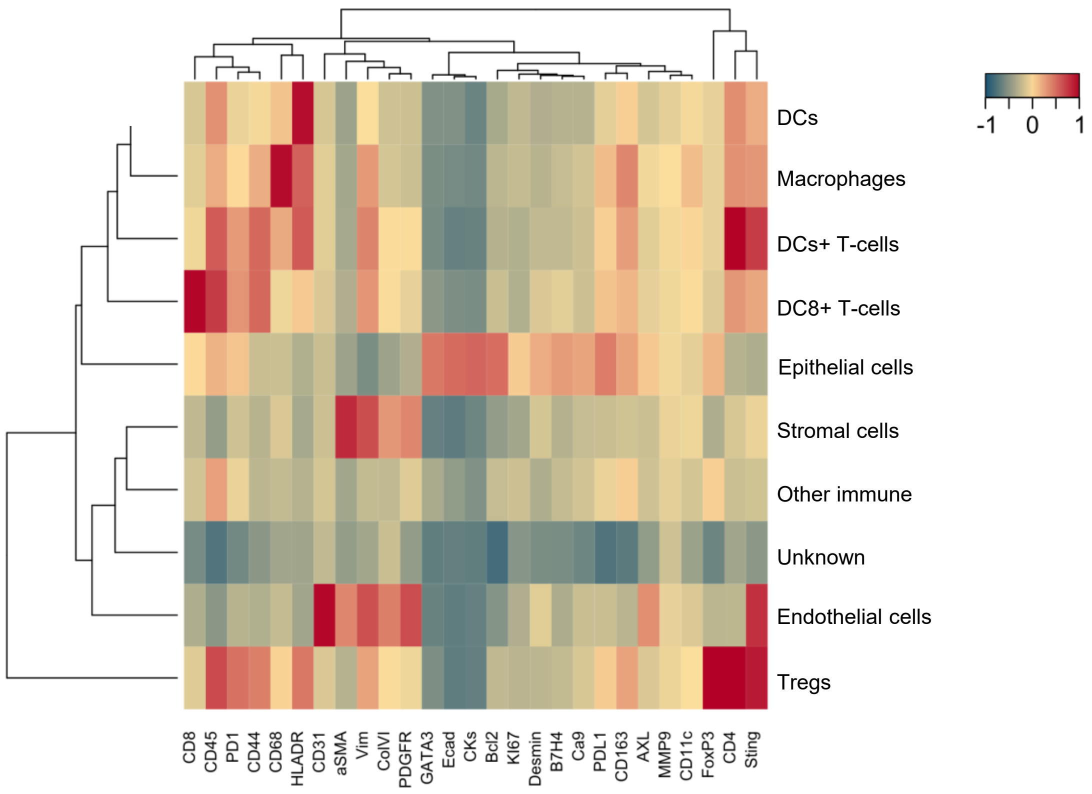


**Supplementary Fig. S2. Cell type protein expression.** Heatmap of mean protein expression in eight cell phenotypes including "unknown" and "other immune" grouped with hierarchical clustering.

**Supplementary Fig. S3: Longitudinal AMD and colocalization score of epithelial cells/target cells**

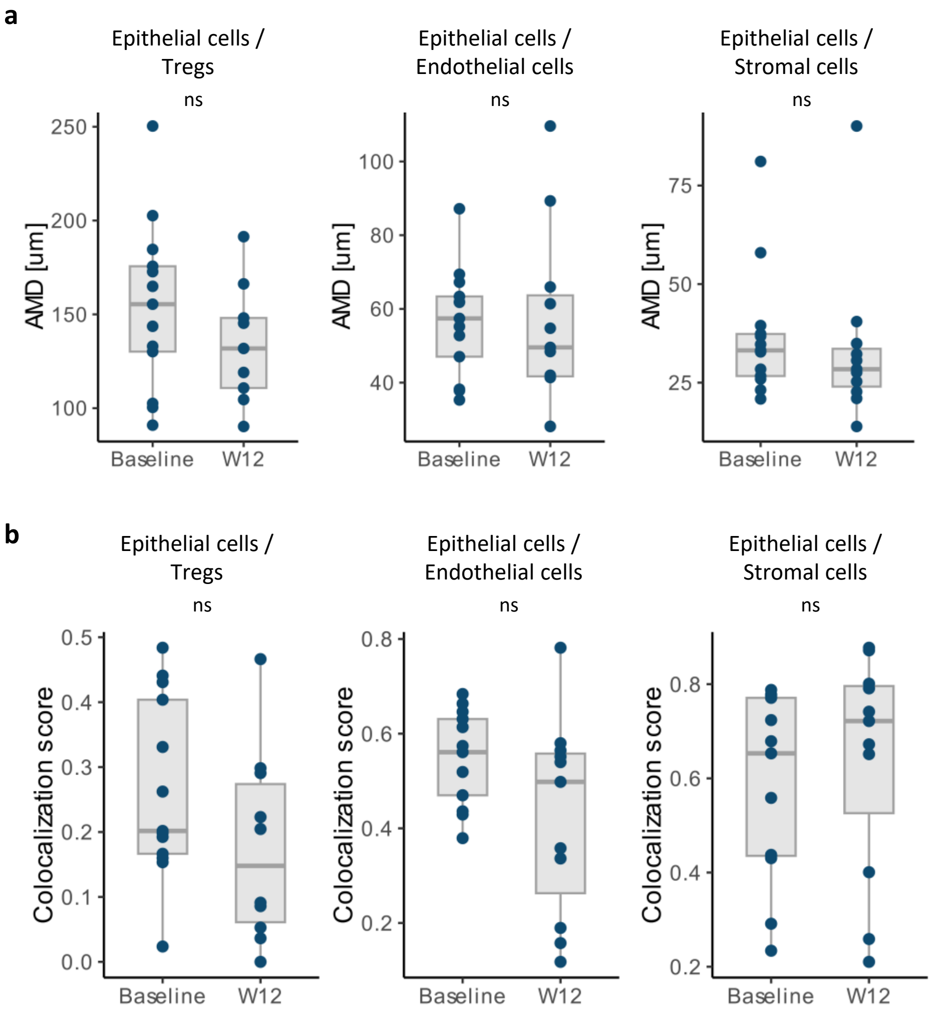


**Supplementary Fig. S3. Longitudinal AMD and colocalization score of epithelial cell/target cell pairs. a,b.** Average minimum distance (AMD) (a) and colocalization score (b) between epithelial cells and Tregs, endothelial cells and stromal cells at baseline and on-treatment (W12). ns: not significant.

**Supplementary Fig. S4: Longitudinal AMD and colocalization score of CD8+ T-cells/target cells**

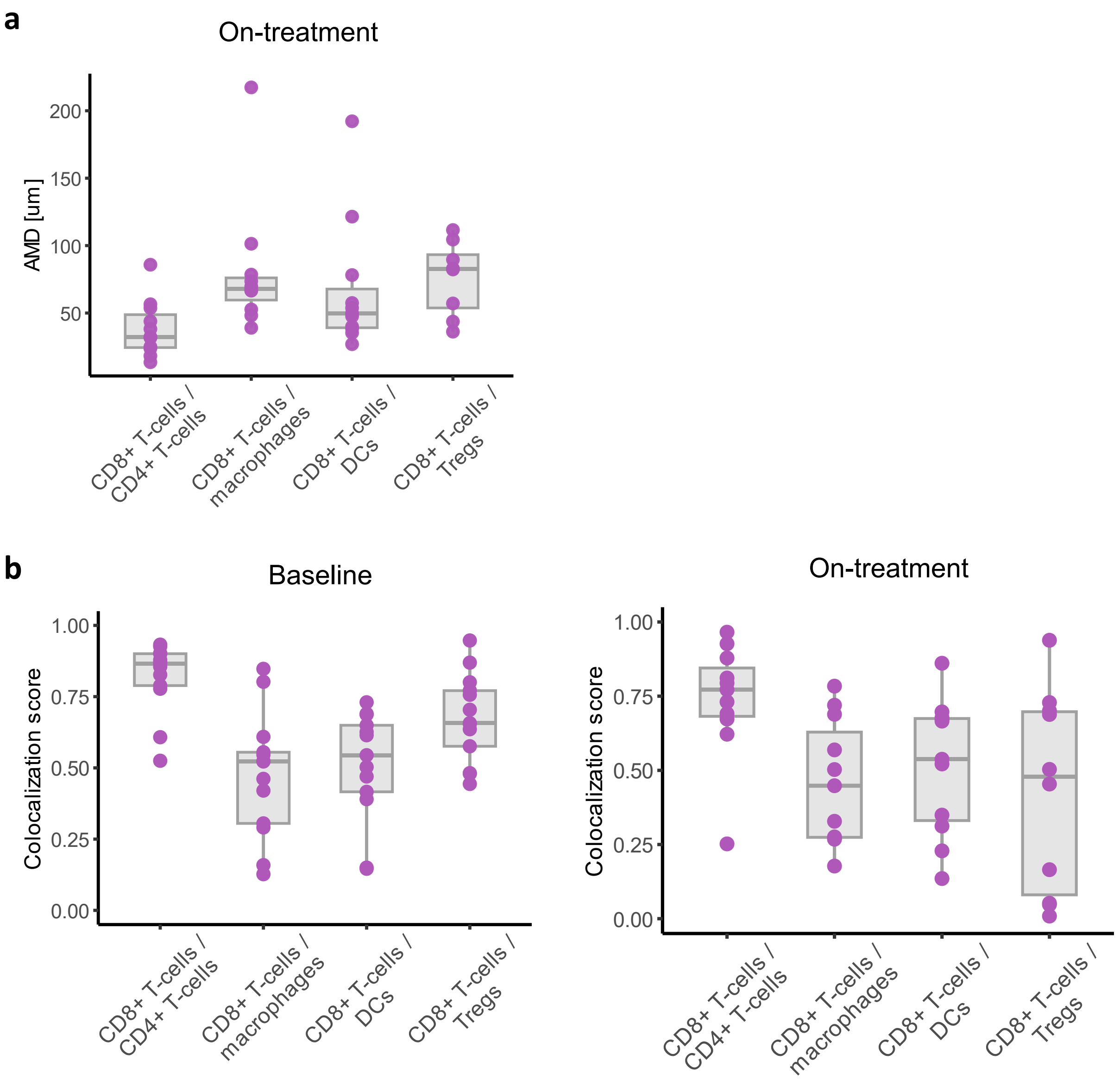


**Supplementary Fig. S4. Longitudinal AMD and colocalization score of CD8+ T-cell/target cell pairs. a,b.** Average minimum distance (AMD) (a) and colocalization score (b) between CD8+ T-cells and CD4+ T-cells, macrophages, dendritic cells and Tregs at baseline and on-treatment (W12).

**Supplementary Fig. S5: Cell density and treatment response**

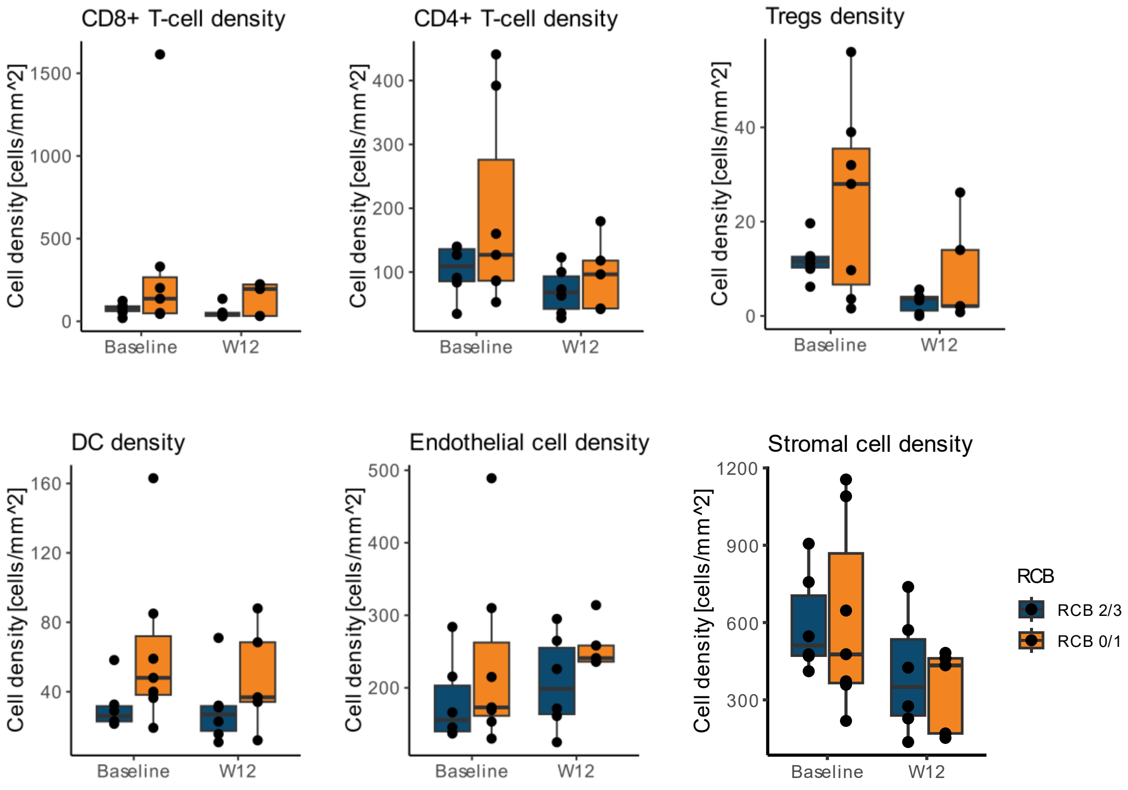


**Supplementary Fig. S5. Cell density and treatment response.** Cell density of CD8+ T-cells, CD4+ T-cells, Tregs, dendritic cells, endothelial cells, and stromal cells separated by treatment response (RCB 0/1 vs. RCB 2/3) at baseline and on-treatment (W12).

**Supplementary Fig. S6: Epithelial cell/target cell AMD and treatment response**

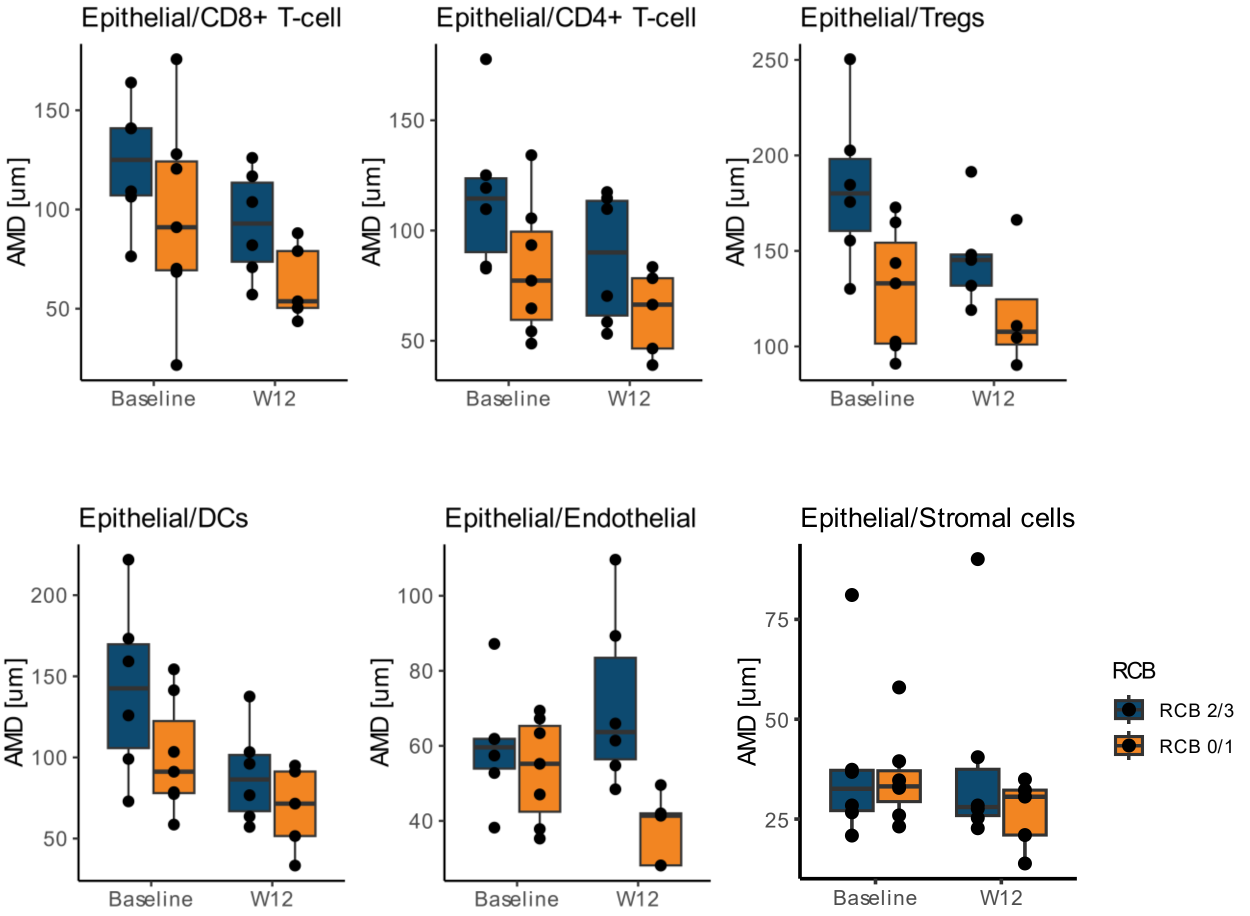


**Supplementary Fig. S6. Epithelial cell/target cell AMD and treatment response.** AMD between epithelial cells and CD8+ T-cells, CD4+ T-cells, Tregs, dendritic cells (DCs), endothelial cells, and stromal cells separated by treatment response (RCB 0/1 vs. RCB 2/3) at baseline and on-treatment (W12).

**Supplementary Fig. S7: Density of CD8+ T-cell subtypes**

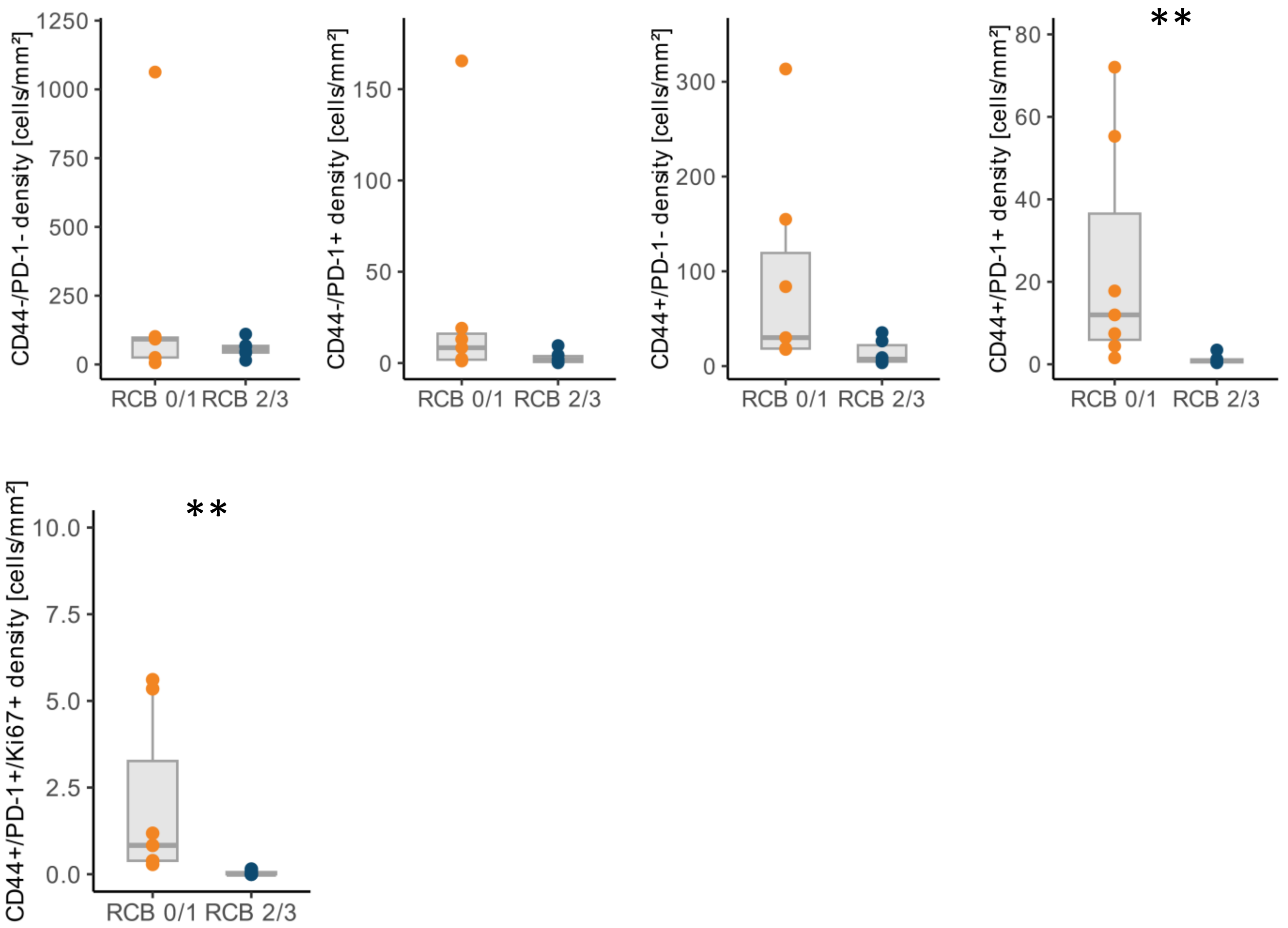


**Supplementary Fig. S7. Density of CD8+ T-cell subtypes.** Density of CD8+ T-cell subtypes separated by treatment response (RCB 0/1 vs. RCB 2/3).

**Supplementary Fig. S8: longitudinal fraction of CD8+ T-cell subtypes**

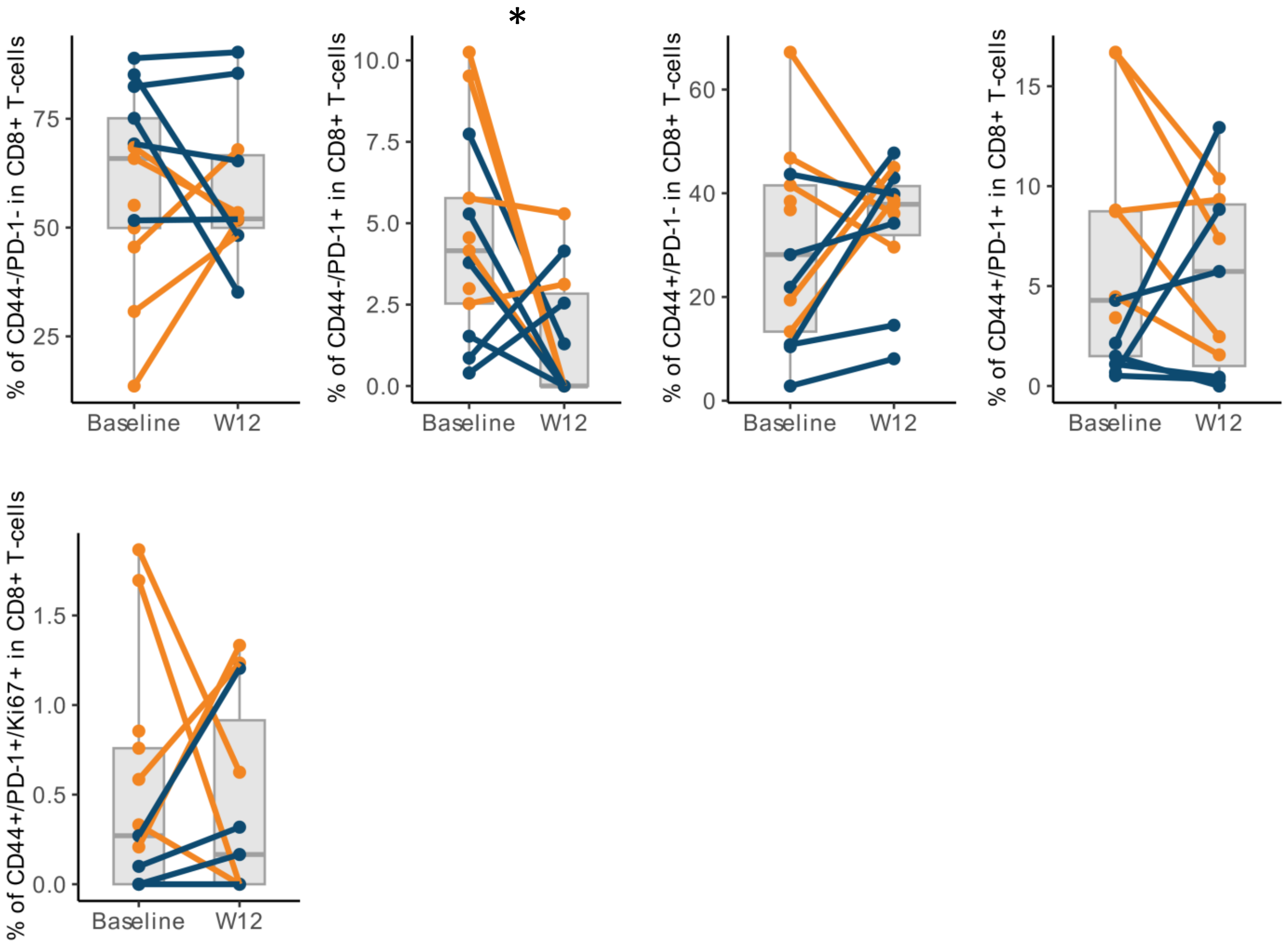


**Supplementary Fig. S8. Longitudinal fraction of CD8+ T-cell subtypes.** Fraction of CD8+ T-cell subtypes in responders (orange) and non-responders (blue) at baseline and on-treatment (W12).
