## Supplementary Methods for "Spatial organization of the tumor-immune microenvironment in ER-positive breast cancer: remodeling during treatment and associations with clinical response"

**Calculation of average minimum distance (AMD)**

The minimum distance defined as the distance between a reference cell and the closest target cell was calculated with vector-based analysis using the Euclidean distance from cell to cell. As a metric of cell proximity, the average minimum distance (AMD) was calculated by averaging the minimum distance for all reference cell/target cell pairs. AMD was defined as

$AMD= \frac{\sum d_{min}^{ref\to target}}{\# refence cells}$,

where $d_{min}^{ref\to target}$ is the distance from a reference cell to the nearest target cell. Epithelial cell interactions were analyzed with epithelial cells as reference cell type and CD8+ T-cell interactions were analyzed with CD8+ T-cells as reference cell type.

We compensated for edge effects by excluding any reference cell for which the closest target cell was found at a distance further than the closest tissue sample outline or defect area in the calculation.

**Calculation of the Morisita colocalization score**

The Morisita similarity index (colocalization score) was calculated as a global measure of degree of colocalization between two cell types and measures how similar two cell type distributions are within the tissue sample. The score was calculated by partitioning the sample image into grid tiles (side length of 250 µm) and calculate the proportional distribution of each cell type within each tile. The score was defined as

$M= \frac{2\sum_{i} p_{i}^{a}p_{i}^{b}}{\sum_{i} \left( p_{i}^{a} \right)^{2}+\sum_{i} \left( p_{i}^{b} \right)^{2}}$,

where $p_{i}^{a}$and $p_{i}^{b}$ are the proportion of cell type *a* and *b*, respectively, within tile *i*. The method generates a score between 0 and 1, where 1 indicates that the cell types are highly colocalized, 0.5 that the cell types are spatially independent, and 0 indicates that the cell types are completely spatially separated.

**Marker normalization for macrophage characterization**

Macrophage subtypes were defined using the markers HLA-DR, CD163, PD-L1 and CD11c. To normalize marker expressions, values were rescaled within each sample using two reference points selected from the marker distributions of macrophages and stromal cells serving as control. Rescaling was performed as:

$$\tilde{m}=\frac{m-p_{1}}{p_{2}-p_{1}}$$

where $\tilde{m}$ is the rescaled marker value, $m$ the unscaled marker value, $p_{1}$ the lower reference point, and $p_{2}$ the upper reference point. Following rescaling, the data from all samples was combined and standardized.

Macrophage subtypes were identified by hierarchical clustering of the selected markers (Euclidean distance, Ward’s method). The optimal number of clusters was determined to be three based on visual analysis of the dendrogram and silhouette score.
