## Supplementary Table S1 for "Spatial organization of the tumor-immune microenvironment in ER-positive breast cancer: remodeling during treatment and associations with clinical response"

| Round | Channel 1 | Channel 2 | Channel 3 | Channel 4 | Channel 5 |
| --- | --- | --- | --- | --- | --- |
| 1 | DAPI | AF488 | AF555 | AF647 | AF750 |
| 2 | DAPI | ColVI | CD31 | CD4 | ECadherin |
| 3 | DAPI | Desmin | B7H4 | CD8 | CD20 |
| 4 | DAPI | aSMA | CD68 | PD-1 | CD45 |
| 5 | DAPI | Vimentin | AXL | PD-L1 | FoxP3 |
| 6 | DAPI |  | Ca9 | CD163 | Ki67 |
| 7 | DAPI | panCK | Bcl2 | CD44 | HLA-DR |
| 8 | DAPI |  | PDGFR | MMP9 | GATA3 |
| 9 | DAPI |  | CD11c | Sting | CD11b |
